# Non canonical activation of the ESCRT machinery is required for division of *Leishmania donovani* parasitophorous vacuoles and parasite persistence

**DOI:** 10.1101/2024.07.06.602309

**Authors:** Javier Rosero, Peter E. Kima

## Abstract

In the mammalian host, *L. donovani* are intracellular pathogens that reside in vacuolar compartments (often called *Leishmania* parasitophorous vacuoles (LdLPVs)). LdLPVs harbor individual parasites that enigmatically divide upon replication of the parasite. In this study, we evaluated the role of the ESCRT machinery in the division of LdLPVs and parasite persistence in infected cells. We found that the ESCRT I member, TSG101 and the ESCRT III members, CHMP2B and CHMP4B are recruited to LdLPVs. In addition, Vps4a, an accessory molecule required for recycling of ESCRT III molecules is also recruited to LdLPVs. Interestingly, infection of cells expressing a dominant negative version of Vps4a that prevents the recycling of ESCRT III revealed that most LdLPVs recruit ESCRT components constitutively. Based on that finding, we proposed that the recruitment of ESCRT molecules to LdLPVs is enabled by the display of the phosphoinositide, PI(3,4)P2 on LdLPVs. To assess the functional importance of recruiting ESCRT molecules to LdLPVs, we monitored *L. donovani* infections in cells in which ALIX or TSG101 were knocked down. ALIX knock down resulted in LdLPVs that were distended and harbored 4 or more parasites, which is significantly different from LdLPVs in ‘wild type’ macrophages that harbor at most, 2 parasites. Moreover, reduced levels of ALIX resulted in a significant reduction in parasite numbers. These findings revealed the critical role for activation of the ALIX-ESCRTIII axis in *L. donovani* pathogenesis. This is the first demonstration that the ESCRT machinery plays a role in the division of pseudo-organelles that harbor an intracellular pathogen.

**Significance:** The endosomal sorting complex required for transport (ESCRT) machinery plays critical mechanistic roles in physiological processes including cell division (cytokinesis). It can be hijacked to promote the spread and persistence of infectious agents including in the budding of viruses and nutrient acquisition by intracellular pathogens. In this study, we uncover a new role for the ESCRT machinery in the infection of macrophages by *Leishmania donovani* (Ld). Within infected cells, each Ld parasite resides in a *Leishmania* parasitophorous vacuole (LPV) that enigmatically divides to accommodate daughter parasites. We show that a non-canonical activation of the ESCRT machinery is required for division of LPVs and for parasite persistence. Future studies on the mechanisms for selective activation of the ESCRT machinery would reveal targets for the control of this deadly pathogen.

## Introduction

*Leishmania* spp. parasites are single celled protozoan organisms that are transmitted by sandflies. *Leishmania donovani* is the causative agent of visceral leishmaniasis that has an estimated 50,000 to 90,000 new cases per year, 95% of which are fatal if left untreated (https://www.who.int/news-room/fact-sheets/detail/leishmaniasishttps). Visceral leishmaniasis presents as irregular bouts of fever, weight loss, enlargement of the spleen and liver, and anemia [1]. In addition to infection of monocytes and macrophages, *L. donovani* infect hematopoietic stem cells, megakaryocytes and other cell lineages that are not known to be phagocytic [2]. Once inside mammalian cells, *Leishmania* spp. are lodged in membrane enclosed compartments often called the *Leishmania* parasitophorous vacuole (LPVs). LPVs fuse extensively with late endosomal compartments and with vesicles from the secretory pathway [3–5]. Interestingly, LPVs that harbor *L. donovani* (LdLPVs) acquire late endosomal characteristics slowly as evidenced by their prolonged retention of Rab5 among other early endosomal molecules [6]. Moreover, LdLPVs harbor individual parasites and enigmatically divide following parasite replication to accommodate daughter parasites that segregate into separate LdLPVs [4,7]. *Leishmania* replication in mammalian cells is asynchronous and commences approximately 24 hours after parasite internalization [8]. The division of LdLPVs stands out as a mechanistic conundrum. As we considered the likely underlying mechanisms of LdLPV division, we wondered whether it is mechanistically like cytokinesis where a cytoplasmic bridge at the mid body between nascent daughter is eventually cleaved in a process called abscission [9,10]. Alternatively, LdLPV division could be mechanistically like the budding of enveloped viruses from membrane delimited compartments including the nuclear or plasma membrane as described in studies on the egress of EBV from the nucleus or HIV from the plasma membrane of infected cells [11,12]. In both cytokinesis and viral budding, the endosomal sorting complex required for transport (ESCRT) machinery has been implicated.

The ESCRT machinery is composed of 4 molecular complexes that are functionally conserved in eukaryotes. In mammalian cells, ESCRT 0 members include the hepatocyte growth factor-regulated tyrosine kinase substrate (HGRS-1) and signal transducing adaptor molecule (STAM) 1 and 2 [13]. The ESCRT I complex includes tumor-susceptibility gene 101 (TSG101), vacuolar protein sorting (VPS) members VPS28, VPS37 (A-D) and UBAP1 [14]. ESCRT II includes subunits of ELL-associated protein 20/vacuolar protein sorting (EAP) (20, 30 and 45) [15]. While ESCRT III includes the Charged membrane proteins (CHMPs) 1, 2, 3, 4, 6 and 7, with 2 variants for CHMP1 (a and b), 2 variants for CHMP2 (a and b) and 3 variants of CHMP4 (a, b, and c) [16]. In addition to these complexes, there are accessory molecules that associate with some complexes and that play crucial roles in the overall functions of the ESCRT machinery. Notably, ALG-2 interacting protein X (ALIX) is an accessory molecule to the ESCRT I complex and may substitute for ESCRT I in some activities [17]. The AAA ATPase vacuolar protein sorting-associated protein (VPS4 (a and b)) is another accessory molecule that interacts most extensively with members of the ESCRT III complex and catalyzes membrane scission by ESCRT III and promotes their recycling from membranes [18]. Activation of the ESCRT machinery for the formation of intraluminal vesicles within multivesicular endosomes, for example, follows a canonical scheme where there is sequential recruitment of ESCRT complexes [19,20]. Increasingly, studies on the function of the ESCRT machinery are revealing non canonical participation of ESCRT complexes [17,21,22] where some complexes appear to be dispensable to accomplish tasks of the ESCRT machinery. For example, although it is still unsettled, some studies have found ALIX to have a more essential role for the completion of cytokinesis [23–25]. Similar observations were made in studies of plasma membrane repair [26] and HIV budding from the plasma membrane [11]. Such non canonical schemes for the activation of the ESCRT machinery has led to approaches that selectively target some ESCRT molecules to limit well defined ESCRT dependent processes [22,27,28]. Conversely, there is the occurrence of lethal diseases due to defects in select ESCRT members, which underscores the important roles that the ESCRT machinery plays in the maintenance of good health [29].

The ESCRT machinery has been implicated in the pathogenesis of intracellular bacteria and parasites that reside within vacuolar compartments in infected cells [30]. Most intracellular bacteria express secretion systems whose components are inserted into the vacuole delimiting membrane. Insertion of the secretion apparatus into the vacuole membrane apparently causes some damage, which then activates the ESCRT machinery that responds to any damage in the endomembrane systems [31,32]. Such is the case with *Mycobacteria* infections where damage of the *Mycobacterium* containing compartment (MCV) has been shown to occur due to the insertion of ESX machinery, components of the Type VII secretion system [33,34]. Damage of the MCV is known to result in spillage of *Mycobacteria* derived molecules including nucleic acids to the cytosol where they activate the cytosolic surveillance response that induces innate immune responses. The ESCRT machinery is then activated to repair the damaged MCV endomembrane [34–37]. In *Salmonella* infections of epithelial cells, where bacteria reside in the *Salmonella* containing compartment (SCV), which is an intricate network of tubules, the bacteria express the Type III secretion apparatus that is deployed to release effectors into the cell cytoplasm. It was shown that components of the ESCRT machinery are recruited to the SCV, presumably to limit damage from the secretion apparatus [38]. In *Coxiella burnetii* infections too, damage to their pathogen containing vacuole was suggested by the recruitment of galectin 3, a known indicator of lysosome damage [39]. This was believed to be the trigger for the recruitment of ESCRT components.

Membrane damage is not the only inducer for the recruitment of ESCRT molecules to pathogen containing vacuolar compartments. In *Toxoplasma* infections where the parasite containing vacuoles (TgPV) are non fusogenic, it was found that components of the ESCRT machinery are recruited to the TgPVM [40,41]. There, they interact with parasite derived molecules including the GRA proteins that are inserted into the TgPVM. As regards to the function of the components of the ESCRT machinery that are recruited to the TgPVM, they were implicated in the acquisition of nutrients from the host cell [40]. This function of the ESCRT machinery is evidently different from the canonical functions of the ESCRT machinery in cytokinesis and extracellular vesicle biogenesis [32]. It is not known what are the underlying mechanisms that are preferentially activated for the ESCRT machinery to accomplish such non canonical functions. In the study by Riviera-Cueves et al [40] the interactions of TSG101 with the TgPVM were found to be more consequential for the uptake of nutrients than the interactions of ALIX with parasite molecules at the TgPVM.

There is evidence that some phosphoinositide species are the membrane anchor to which ESCRT components are recruited [42–44]. It was shown that for the completion of cytokinesis, phosphoinositides serve as critical membrane anchors for ESCRT molecules ([10,43]. In the lens of the eye, for example, the PI(3,4)P_2_ (phosphatidylinositol 3,4-bisphosphate), was identified as the binding partner of the ESCRT II complex member VPS36 [42]. Interestingly, absence or loss of either the lipid or the complex member resulted in impaired cytokinesis.

As discussed earlier, *Leishmania* reside in fusogenic endocytic compartments. Unlike intracellular bacteria, *Leishmania* lack a secretion apparatus that delivers pathogen derived molecules across the LPV to the cell cytosol. It is therefore not known whether in the absence of a vacuole damage inducing machinery, there are other inducers of ESCRT recruitment to the LdLPV. In this study we tracked the recruitment of representative ESCRT molecules to LdLPVs in *L. donovani* infected cells. Pursuant to the observation that members of the ESCRT I and ESCRT III complexes are recruited to LdLPVs, their distribution on LdLPVs over a 72-hour infection course was analyzed. We took advantage of the availability of the dominant negative construct of VPS4 (pEGFP-VPS4-E228Q) whose expression has been shown to slow down the recycling of ESCRT III from membranes [40,45], which therefore permits the visualization of transient associations of the ESCRT machinery with membranes. To gain insight on how cells would respond to damage of LdLPVs we studied the effects of the lysosome membrane-rupturing agent l-leucyl-l-leucine methyl ester (LLOME) on the recruitment of ESCRT molecules to LdLPVs. To commence addressing what are the critical roles of an activated ESCRT machinery in *Leishmania* infected cells, we monitored the effect knocking down TSG101 or ALIX. TSG101 and ALIX can function in the recruitment of downstream components including ESCRT III members [21]. The effect of ALIX knockdown was dramatic. There was an increase in the frequency of larger LdLPVs harboring greater than 4 parasites. LdLPVs division was defective in ALIX knockdowns due to failure of the proper activation of the ESCRT machinery. Finally, there was a reduction in overall parasite burden in the cultures, which was likely due to the inability of parasites to complete replication.

## Results

### ESCRT I and ESCRT III molecules are recruited to LdLPVs

ESCRT molecules are cytosolic proteins that are recruited to membranes, where they exercise their functional activities [22,31]. We wanted to know whether the ESCRT machinery is assembled on LdLPVs. To supplement observations using commercially available antibodies to mouse ESCRT molecules, we elected to evaluate the redistribution of fluorophore-tagged recombinants of a representative ESCRT I member, TSG101 (mEGFP-TSG101), representative ESCRT III members, CHMP2B (mCherry-CHMP2B) and CHMP4B (mCherry-CHMP4B), in *L. donovani*-infected cells. We also evaluated the distribution in infected cells, of the ATPase VPS4A (mCherry-VPS4A), which is an accessory molecule that is critical for the recycling of ESCRT III molecules from membranes. On average just under 60% of macrophages are infected under our experimental conditions. Unlike in uninfected cells where there is diffuse punctate labeling of these molecules, in some infected cells, these molecules were recruited to and displayed on LdLPVs. Representative images of the distribution of the tagged molecules in uninfected cells and their recruitment to LdLPVs is shown in Figure 1a. To visualize the contours of the LdLPV membrane, lysosome associated membrane protein 1 (Lamp1) was labeled with either a red or green fluorophore dictated by the fluorophore of the tagged ESCRT member.

**Figure 1.**
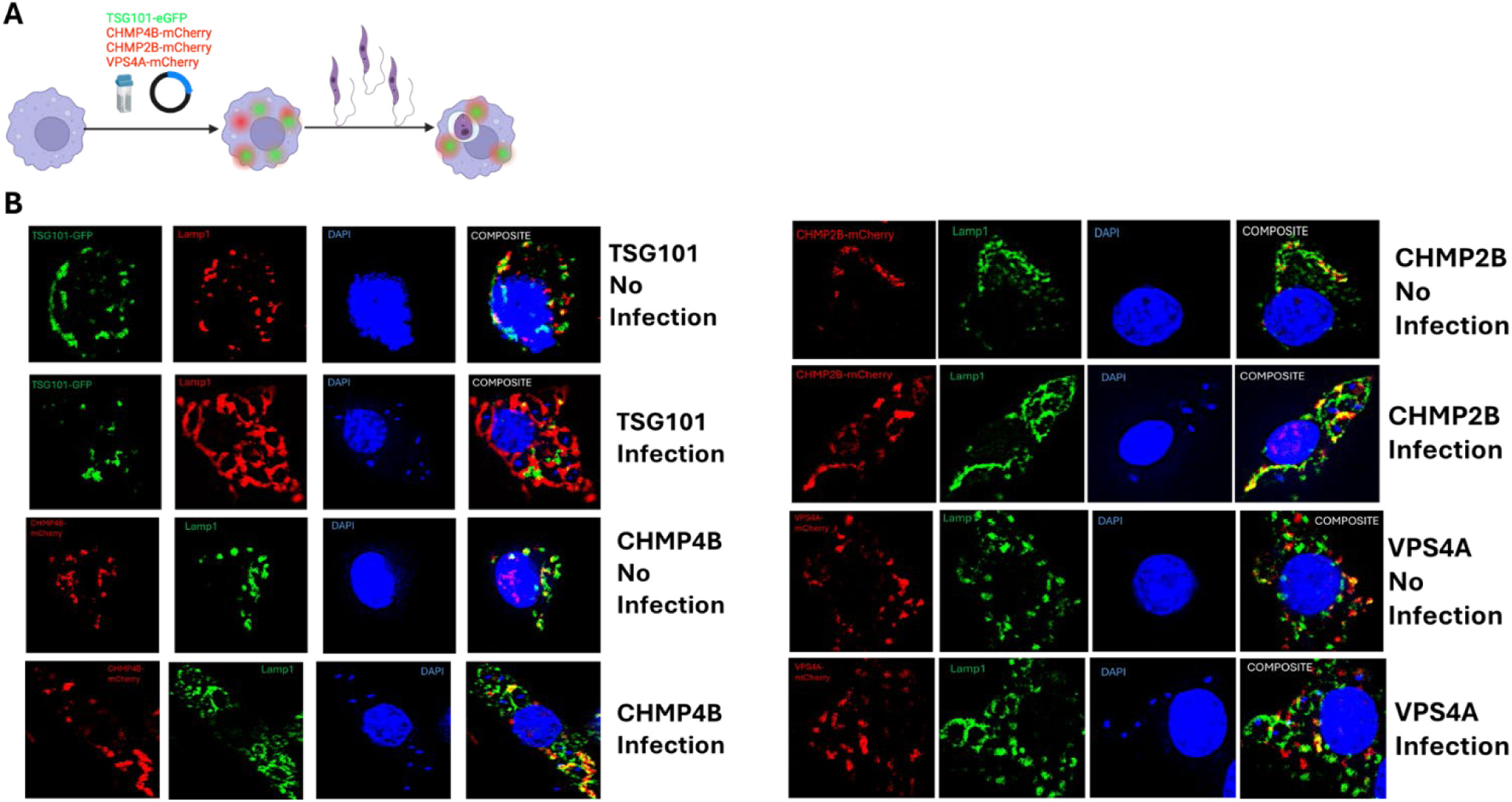
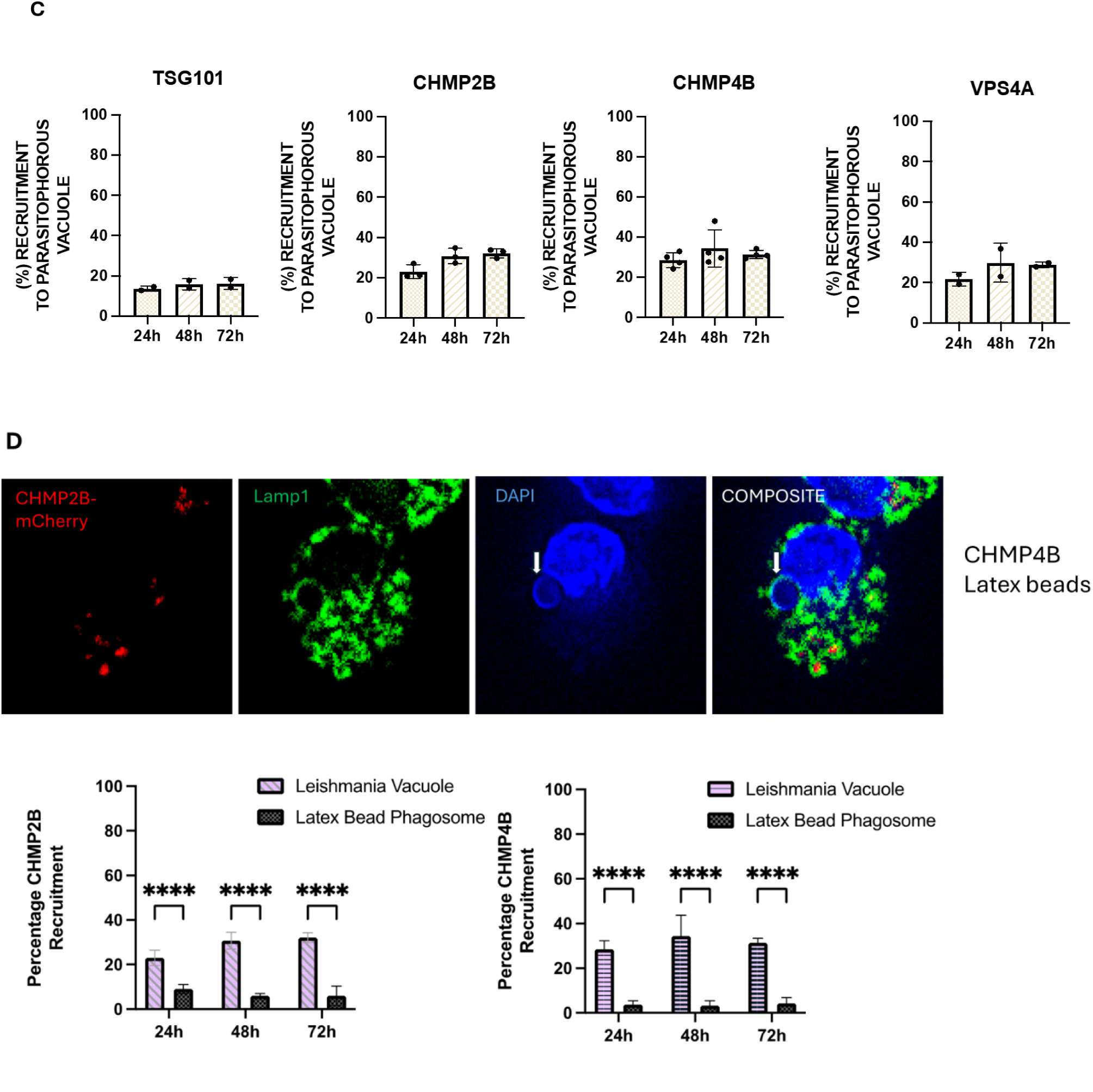
Components of the ESCRT machinery are recruited to LdLPVs Recombinant fluorophore-tagged components of the ESCRT machinery were transiently expressed in RAW264.7 macrophages plated on coverslips. The cells were infected for 24, 48 or 72 hrs with metacyclic *L. donovani* parasites. The color of each fluorophore-tagged recombinant is shown (A). Lamp1 is labeled with either a red or green tagged fluorophore in contrast fluorophore of the recombinant-tagged molecule. Parasite and host cell nuclei are labeled with DAPI (blue). Representative uninfected and infected cells are shown (B). The proportion of Leishmania parasitophorous vacuoles in transfected cells that were deemed to be positively displaying the indicated recombinant fluorophore was counted and plotted (C). Recruitment of ESCRT components to generic phagosomes was determined by incubating transfected cells with latex beads. A representative image of a cell transfected with CHMP4b is shown (D). The proportion of phagosomes that displayed CHMP2b or CHMP4b is shown in the associated graphs. Lamp1 reactivity was used to delineate the vacuole of the parasite or latex bead phagosome. White arrow points to a latex bead. At least 100 parasite vacuoles were counted per coverslip, per timepoint. Counts were done in duplicate coverslips for at least 3 independent experiments. Graphs were generated in GraphPad Prism*8 followed by statistical tests. One-way ANOVA with multiple comparisons followed by *post hoc* Tukey’s honest significant difference test to determine statistical significance was performed. *p<0.05, **p<0.01, ***p<0.001, ****p<0.0001.

The distribution of each of these molecules on LdLPVs followed similar patterns. In come infected cells, the tagged ESCRT molecule was at the midpoint of two LdLPVs. For example, in TSG101-GFP transfected cells, TSG101 is at the midpoint of two LdLPVs (Figure 1b). Images of CHMP4b show labeling at the periphery of the LdLPVs (Figure 1b). CHMP4b was also found in the middle between two LdLPVs. A representative image is shown in Supplementary Figure (Supplementary Figure1). CHMP2b labeled similarly. I representative image is shows of CHMP2b labeling internal of the Lamp1 label. This later labeling pattern is suggestive of a horn-like pattern structure where the neck of the horn is constricted (CHMP2b and CHMP4b label here) but the outer rim is wider (Lamp 1 labels here). Such labeling suggests either vesicles budding from the LPV or the formation of a constricted mid-region during LdLPV fission and scission.

We proceeded to count the number of LdLPVs within transfected cells that were deemed to be positive for each of these molecules. The counts compiled from 3 experiments are shown in Figure 1c. Approximately 15% of LdLPVs displayed TSG101-GFP. Approximately 30% of LdLPVs positively displayed CHMP2B-mCherry or CHMP4B-mCherry by 48 and 72hrs. A comparable number of LdLPVs were positive for VPS4A-mCherry at the same times post infection.

There is presently no information that can provide insight into the expected frequency of the recruitment of ESCRT components to LdLPVs. Rivera-Cuevas et al [40] and others [45,46] had reported on the value of expressing VSP4-E228Q, a dominant negative variant of VPS4A in cells. The inability of this variant to hydrolyze ATP prevents the recycling of ESCRT III complexes from membranes thereby forcing their accumulation. We proceeded to perform a dual transfection of pEGFP-VSP4-E228Q and CHMP4b-mCherry followed by infection with *L. donovani* (Figure 2a). Representative images (Figure 2b) show a more accentuated labeling of CHMP4B-mCherry on LdLPVs. Enumeration of the number of LdLPVs that were positive for CHPMP4B revealed that greater than 90% were positive for this molecule. Evidently when the recycling of ESCRT III molecules was prevented, due to expression of VSP4-E228Q there was an accumulation of CHMP4b on most LdLPVs. This result suggested that the counts for the proportion of LdLPVs with recruited ESCRT molecules presented in (Figure 1c) was possibly an under count of the number of LdLPVs that recruit CHMP4b. Taken together, an interpretation of these results is that some ESCRT molecules are constitutively recruited to LdLPVs. However, a subsequent signal may be required to trigger the aggregation of the ESCRT machinery, which produces more outstanding labeling of ESCRT components.

**Figure 2.**
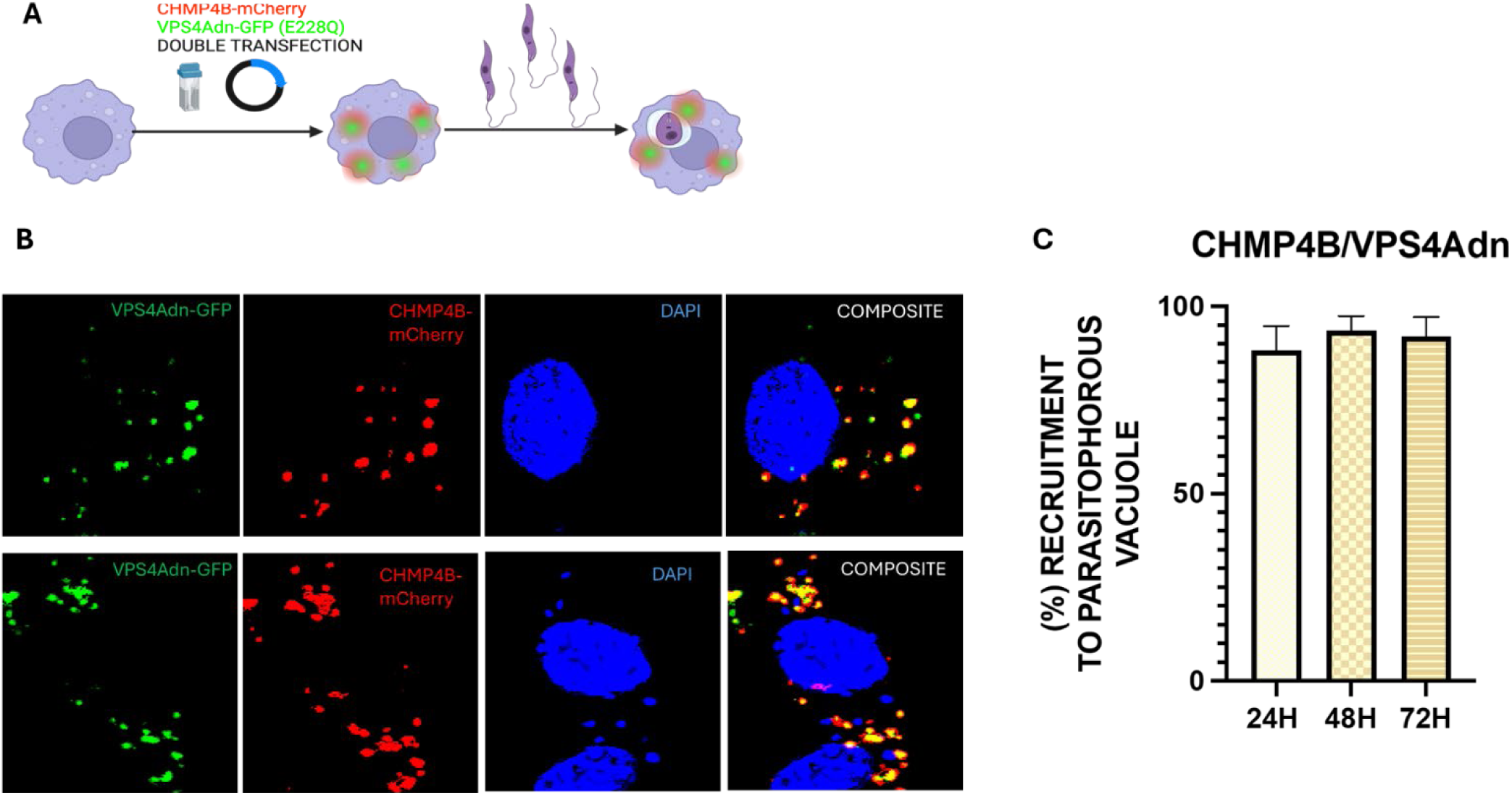
Expression of dominant negative VPS4a (VSP4-E228Q) reveals high recruitment of CHMP4b. Macrophages were transfected with pEGFP-VSP4-E228Q and CHMP4B-mCherry and plated on coverslips. After infection with *L. donovani*, cells on coverslips with fixed at 24, 48 and 72hrs. The experimental scheme is shown in (A). Representative images of and uninfected and infected cell are shown (B). Most vacuoles that were positive for CHMP4b were also positive for VSP4-E228Q (C). Each parasite was assumed to be in its own vacuole, which was then scored for VPS4 reactivity. Counts were done in duplicate coverslips for at least 3 independent experiments. Graphs were generated in GraphPad Prism*8.

Inert particles including latex beads or *Zymosan* particles that are internalized by phagocytosis are often used to establish the interactions of phagocytic compartments as they traverse through the endocytic pathway [47,48]. Here we monitored the recruitment of CHMP2b and CHMP4b to phagosomes harboring 2 micron-sized latex beads. Representative images from CHMPb-mCherry transfected cells harboring latex beads show no recruitment to the phagosomes (images from CHMP2b-mCherry transfection are not shown). The enumeration of phagosomes harboring latex beads (Figure 1d) shows that less than 2% of those vacuoles displayed the ESCRT III molecules. Our interpretation therefore is that the composition and characteristics of LdLPVs that permit the recruitment of ESCRT molecules, is different from the composition and characteristics of generic phagosomes.

### Effect of LLOMe on ESCRT recruitment to LdLPVs

Several studies have shown that damage to lysosomes is a potent activator of the ESCRT machinery [39]. Induced lysosomal damage can be accomplished by treating cells with the lysosomotropic compound L-Leucyl-L-leucine methyl ester (LLOMe) that disrupts the membrane of lysosomes [49]. We sought to determine whether treatment of infected cells with LLOMe would cause damage to LdLPVs, which will induce the recruitment of ESCRT components and the activation of the ESCRT machinery. The scheme for the experiments with LLOMe is shown in Figure 3a. CHMP4b-mCherry or CHMP2b-mCherry transfected cells plated on coverslips were infected with *L. donovani* for 48 hours. Thereafter, the cultures were pulsed with LLOMe (1mM or 5mM) or vehicle for 30 minutes. The cultures were washed and fixed at the indicated times. Almost all lysosomes within treated cells recruited CHMP4b-mCherry or CHMP2b-mCherry, which was monitored here as the positive indicator of LLOMe induced damage. Representative images show that CHMP4b-mCherry is recruited to LdLPVs (Figure 3b). Approximately 95% of LdLPVs recruited CHMP4b-mCherry immediately after the pulse of 5mM LLOMe (Figure 3 C,D) The proportion of positive LdLPVs diminished rapidly when the drug was chased out. There was a significantly lower percentage of LdLPVs that recruited CHMP2b-mCherry after 5mM LLOMe treatment as compared to CHMP4b (Figure 3 E,F). This observation suggested that ESCRT III components are differentially recruited to LdLPVs to repair induced damage. This pattern of differential recruitment of CHMP2b and CHMP4b contrasts with the observations presented earlier on the recruitment of these ESCRT III molecules to LdLPVs. Although there is the possibility that damage to LdLPVs may occur, which will necessitate repair, the LLOMe experiments suggests that recruitment for that purpose may be mechanistically distinct from the ‘constitutive’ recruitment of ESCRT molecules to LdLPVs, where CHMP2b and CHMP4b are recruited equivalently (Figure 1).

**Figure 3.**
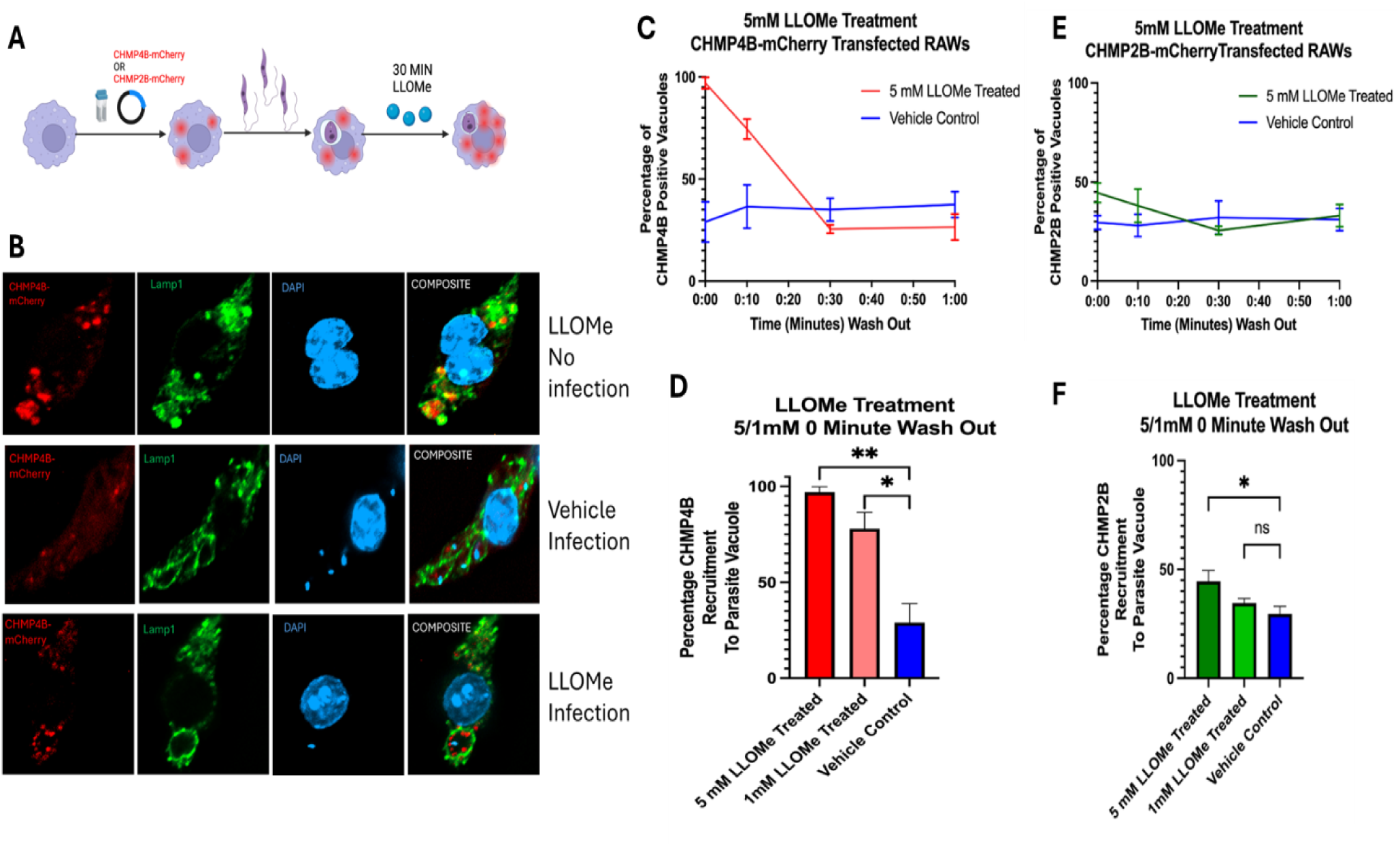
LLOMe induced damage of LdLPV suggests that damage dependent ESCRT recruitment is mechanistically different from ‘normal’ scheme of ESCRT recruitment to LdLPVs. The scheme for the experiments is shown (A). Cells were transfected with either CHMP2b-mCherry or CHMP4b-mCherry and plated on coverslips. They were then infected for 48 hrs after which they were incubated with LLOMe (5mM) or vehicle for 30 minutes. Cover slips were recovered when drug was washed out (time 0) and then subsequently after 10 minutes intervals. Representative images of LLOMe treated CHMP4b transfected cells without infection are shown (red circles are due to CHMP4b in damaged lysosomes) (B). Images from CHMP4b transfected and infected cells treated with vehicle or with the drug are shown. Lamp1 labeling was used to delineate the contours of LPVs. The number of LPVs that were positive for CHMP4b after the pulse and chase of LLOMe is shown (C). A graph of the proportion of positive cells at the 0 time is shown (D). The number of LPVs that were positive for CHMP2b after the pulse and chase of LLOMe is shown (E). A graph of the proportion of CHMP2b positive cells at the 0 time is shown (F). These graphs were compiled from two experiments with 3 coverslips for each point. Graphs were generated in GraphPad Prism*8 and statistical analysis was performed. One-way ANOVA with multiple comparisons followed by *post hoc* Tukey’s honest significant difference test to determine statistical significance was performed. *p<0.05, **p<0.01, ***p<0.001, ****p<0.0001.

### The phosphoinositide PI(3,4)P2 on LdLPVs may be the target for recruitment of ESCRT molecules

The observation that ESCRT molecules are constitutively and preferentially recruited to LdLPVs as compared to phagosomes that harbor latex beads, prompted us to consider differences in the molecular composition of LdLPVs that might play a role in interactions with ESCRT components. Several studies have shown that some phosphoinositides displayed on endomembranes are preferred membrane binding anchors for the recruitment of ESCRT proteins. Specifically, the phosphoinositide PI(3,4)P2 has been shown to bind to ESCRT II, which recruits the ESCRT machinery that is indispensable for cytokinesis in the eye [42,50]. In our previous studies, we had shown that LPVs that harbor either *L. amazonensis* or *L. donovani* parasites display PI(3,4)P2, which was in contrast to phagosomes that harbored *Zymosan* particles that do not display this lipid [48]. In the studies here, we sought to affirm that LdLPVs display PI(3,4)P2. Infected cells were fixed and processed for the visualization of PI(3,4)P2. The representative images show labeling of PI(3,4)P2 on LdLPVs (Figure 4). We proceeded to treat infected cells with LY294002 (50μM) which is known to block the synthesis of most phosphoinositides [51]. After treatment of infected cells for 8hrs the display of PI(3,4)P2 on LdLPVs was inhibited. Moreover, the recruitment of CHPM4b was disrupted by treatment with the drug (Figure 4b). There was some increase in the frequency of LdLPVs that harbored more than 2 parasites (not shown).

**Figure 4.**
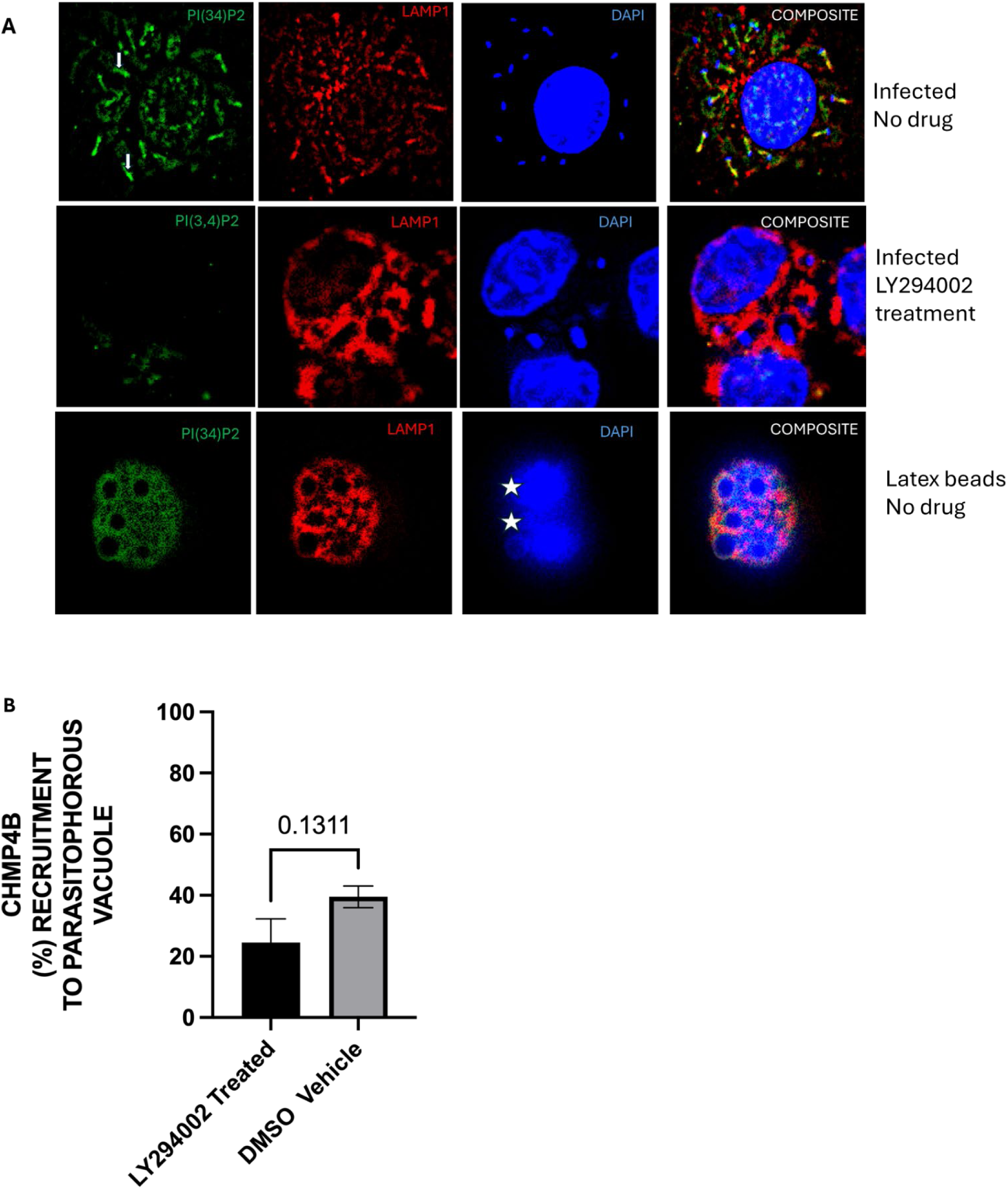
PI(3,4)P2 is displayed on LdLPVs but not on latex bead phagosomes. LY294002 treatment inhibits PI(3,4)P2 display on LdLPVs. Macrophages were infected with *L. donovani*. Representative image of a 48 hrs infected macrophage labeled for detection of PI(3,4)P2 and Lamp1 is shown. This image was selected from more than 3 experiments. Cells infected for 24 hours were incubated in 50uM LY294002 and coverslips were analyzed after 8 hrs. Image is representative from 2 experiments. Latex beads were fed to macrophages. After 24 hrs cells on coverslips were analyzed for the display of PI(3,4)P2 and Lamp1. This image is representative from 2 experiments (A). Cells were transfected with CHMP4b-mCherry, plated and infected. They were then incubated in 50uM LY294002 or vehicle and coverslips were analyzed after 8 hrs for recruitment of CHMP4b to LPVs. At least 100 LPVs were counted per cover slip. These graphs were compiled from two experiments with 3 coverslips for each point. Graphs were generated in GraphPad Prism*8 where unpaired student T-test statistical analysis was performed. *p<0.05, **p<0.01, ***p<0.001, ****p<0.0001.

### Knock down of ALIX disrupts the recruitment of CHMP4B to LdLPVs

What is the functional role of the ESCRT molecules that are recruited to LdLPVs? Several studies have described non canonical schemes for the activation of the ESCRT machinery. In cytokinesis, for example, studies have shown that ESCRT activation follows parallel schemes that are dependent on either the recruitment of ALIX or the recruitment of TSG101, which is followed by the recruitment of ESCRT III components in both schemes [21]. Studies on the budding of HIV from infected cells have also shown greater dependence on the availability of ALIX as compared to TSG101 [52]. Informed by such observations, we proceeded to implement an experimental scheme in which we knocked down either ALIX or TSG101 then monitored several parameters including the recruitment of CHMP4b to LdLPVs, the division of LdLPVs and parasite burden of infected cells (Figures 5a and 6a). Knock downs (KDs) were accomplished by transfection of shRNA plasmids into RAW264.7 cells followed by selection in puromycin and evaluation of oligoclonal lines by Western blotting. Figure 5b shows representative Western blots in which TSG101 or ALIX was detected in the lysates from shCTRL or shTSG101 or shALIX. Oligoclonal lines of ALIX-KD and TSG101-KD with greater than 60% knock down were selected (Figure 5b). We also selected cells transfected with shRNA control plasmid (shCTRL). Knocking down these molecules was not found to affect macrophage viability as measured in an LDH cytotoxicity assay and an MTT proliferation assay (Supplemental Figures 2 a and b).

**Figure 5.**
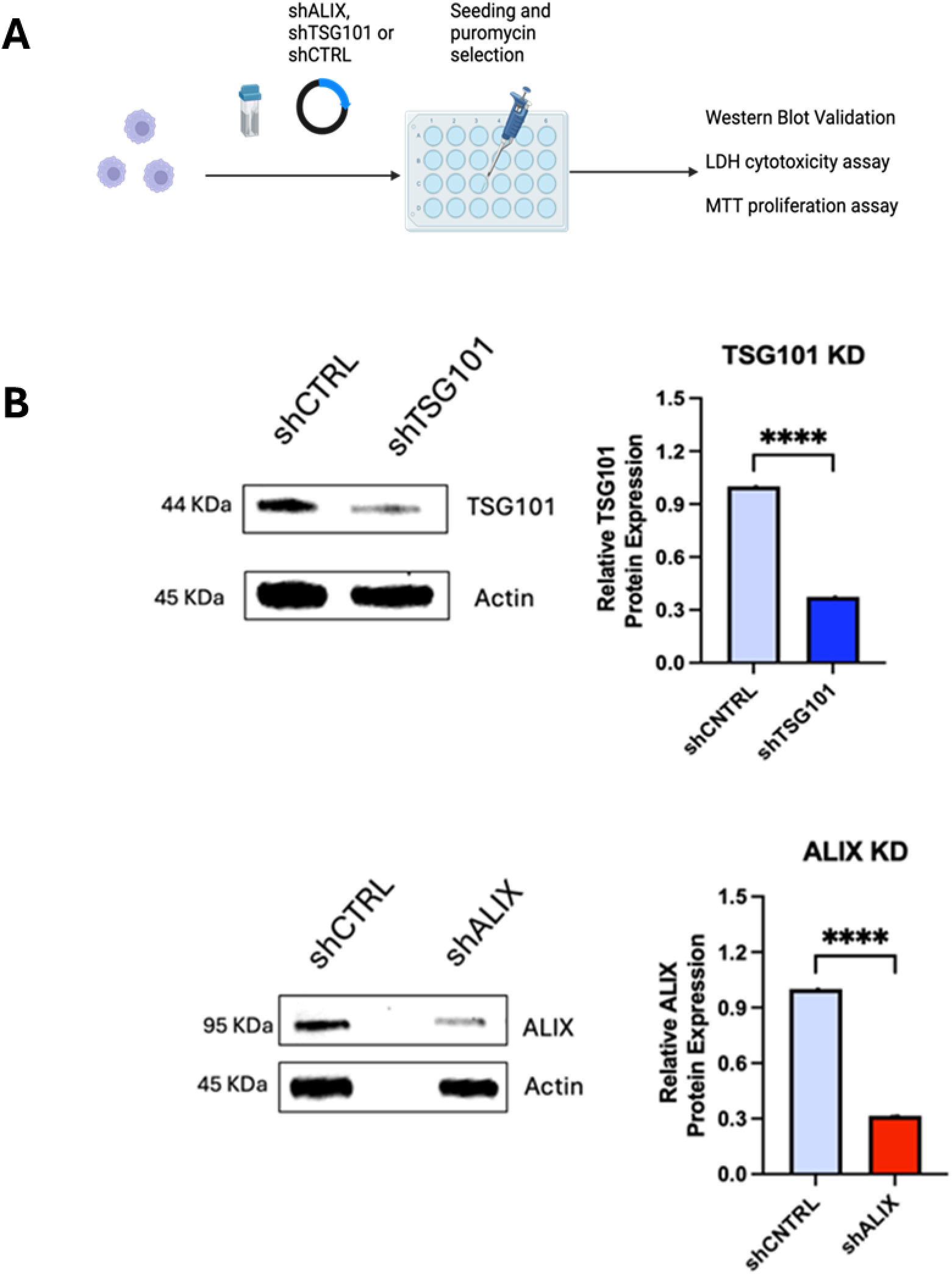
Generation of cells with knocked down levels of TSG101 or ALIX The recombinant TSG101 shRNA plasmid, Alix shRNA plasmid and Control shRNA plasmid were transfected into macrophages then plated. 24 hrs after plating, selection of cells by growth in puromycin was commenced. Oligoclonal cell lines were identified and the scheme for testing of cell lines is shown (A). Oligoclonal lines were tested for the expression of TSG101 or ALIX by Western blot. Representative blot and densitometric analysis of blots for expression of these molecules is shown in (B). Plots were from densitometry of blots from at least 2 experiments. Graphs were generated in GraphPad Prism*8 where unpaired student T-test statistical analysis was performed. *p<0.05, **p<0.01, ***p<0.001, ****p<0.0001.

**Figure 6.**
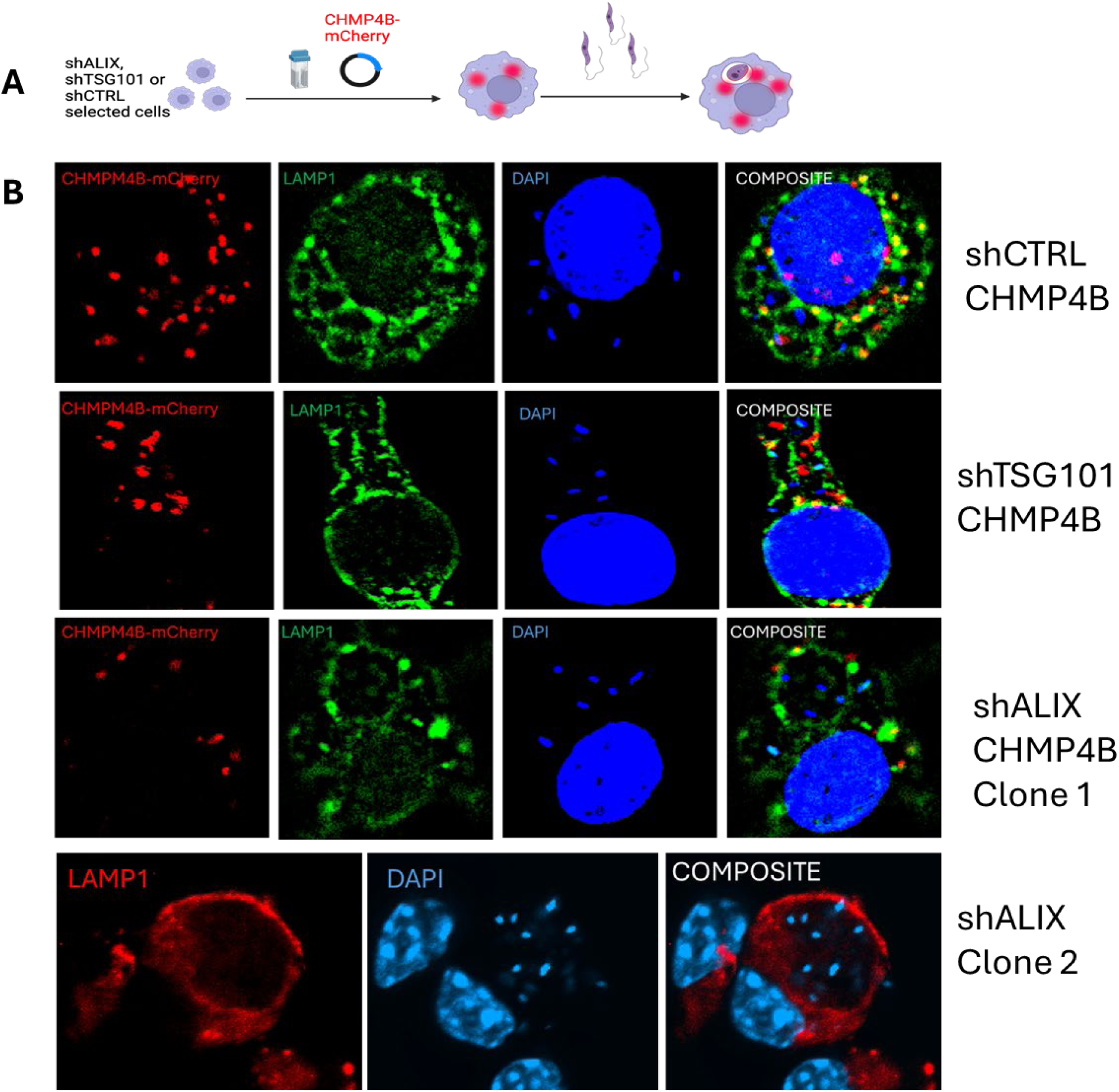
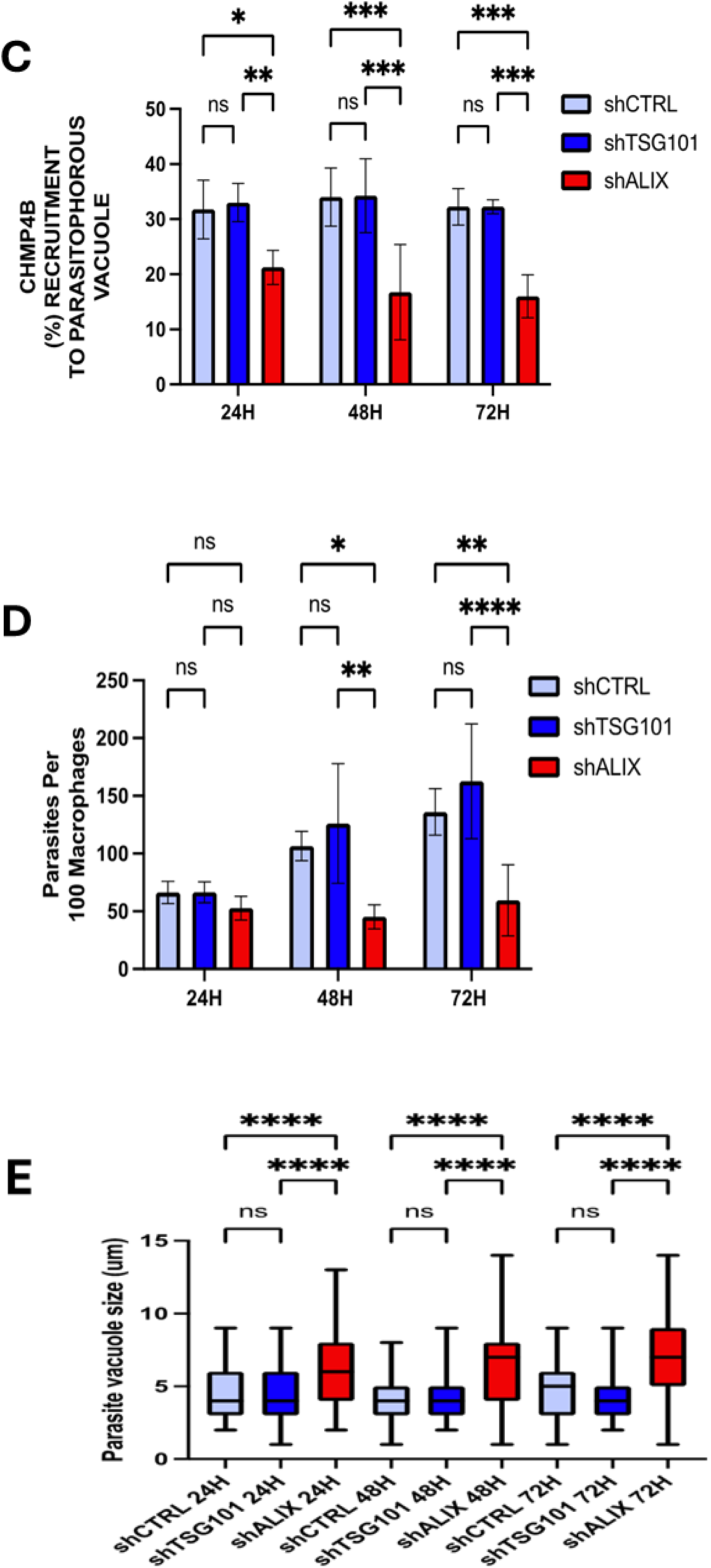
Availability of ALIX as compared to TSG101 is essential for LdLPV division and parasite replication in infected cells. Oligoclonal knockdown cell lines including control knockdown line, were transfected with CHMP4b-mCherry and plated. They were then infected with *L. donovani* parasites as shown in scheme (A). Coverslips were recovered after 24, 48 or 72hs infection and then processed for detection CHMP4b recruitment to LdLPV, Contours of LdLPV membrane was detected by Lamp1 labeling. The parasite and host cell nuclei were labeled with DAPI. Representative images from representative knockdown cell lines, shCTRL, shTSG101 or shALIX are shown (B). shALIX clone 2 was not transfected with CHMP4b-mCherry. (C) The LPV size (parasite vacuole size) in oligoclonal knockdown cell lines after 24, 48 and 72 hrs infection was measured and plotted. Using the “Insert” scale tool of the BZ-X800 analyzer, diameter of parasite vacuoles was measured. At least 100 parasite vacuole sizes were measured per coverslip. Data was compiled and graphed using GraphPad Prism 8. A one-way ANOVA was performed, and statistical significance was determined by *post hoc* Tukey’s honest significant difference test (D) The number of parasites per macrophage in infected oligoclonal knockdown cell lines after 24, 48 or 72hrs infection was enumerated. At least 100 macrophages were counted, per coverslip, per treatment and timepoint. Data was compiled and graphed using GraphPad Prism 8. (E) CHMP4b recruitment to LdLPVs in oligoclonal knockdown cell lines was quantified. Lamp1 reactivity was used to delineate the LPV and colocalization of Lamp1 with CHMP4b was determined. At least 100 LPVs were measured per coverslip, per treatment and timepoint. Data was compiled and graphed using GraphPad Prism 8. One-way ANOVA with multiple comparisons followed by *post hoc* Tukey’s honest significant difference test to determine statistical significance was performed. *p<0.05, **p<0.01, ***p<0.001, ****p<0.0001.

### The knockdown of ALIX, but not TSG101 leads to larger LdLPV that harbor more parasites per vacuole

We then transfected KDs and control lines with CHMP4b-mCherry plasmids, plated overnight and infected with Ld parasites. The scheme for these experiments is shown in Figure 6A. Representative images from the CHMP4b-mCherry transfections in KD cells are shown in Figure 6B. CHMP4b-mCherry was recruited to LdLPVs in shCTRL cells to similar levels as in ‘wild type’ infected cells. Similar proportion of LdLPVs recruited CHMP4b-mCherry in infected TSG101KDs (Figure 6C). In contrast, CHMP4b-mCherry remained diffusely expressed in infected ALIXKDs, with limited recruitment to LdLPVs (Figure 6B,C). Enumeration of the number of LdLPVs that recruited CHMP4B in shCTRL, TSG101KDs and ALIXKDs is shown in Figure 6C. While approximately 33% of LdLPVs in shCTRL and in TSG101KDs were CHMP4b positive at all times post infection, less than18% LdLPVs in ALIXKDs were positive for CHMP4b. The number of LdLPVs that were CHMP4b positive was significantly lower at 24, 48 and 72 hrs post infection.

An often-used indicator of *Leishmania* viability in infected cells is the demonstration that the numbers of parasites per macrophage increases over time, which reflects the replication of viable parasites. In contrast to infection in shCTRL and TSG101KDs where there was the expected increase in the number of parasites/macrophages at 48 and 72 hrs post infection, there was significantly lower parasites/macrophage in ALIXKDs (Figure 6D). Finally, we estimated the sizes of LdLPVs (Figure 6E). The average size of LdLPVs in shCTRL cells was 3.5 microns. In TSG101KDs, LdLPVs were also 3.5 microns. In contrast, the average size of LdLPVs in ALIXKD was approximately 6.8 microns on average, which was significantly higher than LdLPVs in shCTRL and TSG101KDs. As the plots show, in ALIXKDs, there is an increase in the proportion of much larger LdLPVs. Such larger LdLPVs harbored more than 4 −10 parasites. Representative images of large LdLPVs in ALIXKDs that harbor numerous parasites are shown in Figure 6B. We elected to show Images from 2 ALIXKD lines generated from different transfections. These results, including the images show that with reduced availability of ALIX, LdLPV division is defective, which results in approximately 70% reduction in the parasite burden in infected cells.

## Discussion

*Leishmania donovani* are the causative agents of visceral leishmaniasis, which is fatal if not treated. In mammalian cells, Ld are housed individually within membrane enclosed compartments that enigmatically divide to accommodate each daughter parasite after parasite replication. In this study we showed for the first time that components of the ESCRT machinery are recruited to LdLPVs. Surprisingly, reduced availability of ALIX resulted in impaired recruitment of ESCRT III components to LdLPVs. Under those conditions, along with an increase in LdLPV sizes, there was a concomitant increase in vacuoles that harbored multiple parasites. Moreover, there was a significant reduction in the number of parasites in the infected cell culture. Together, these results showed that LdLPV division was impaired due to the loss of ALIX. Evidently, Ld proliferation in infected cells and their overall persistence is dependent on a non-canonical activation of the ESCRT machinery. That a non-canonical activation of the ESCRT machinery plays a role in the division of LdLPVs is a new function for the ESCRT machinery.

To gain more insight into the recruitment of ESCRT components to LdLPVs, we incubated infected cells in LLOMe. This compound is a potent lysosomotropic compound that disrupts the membrane of lysosomes. Several studies have investigated the consequence of exposing cells to this compound [53,54]. We observed that a 30-minute incubation with LLOMe caused rapid damage to most lysosomes, which we detected by the recruitment CHMP4B-mCherry and CHMP2B-mCherry into these compartments. Although most LdLPVs in treated cells became CHMP4B-mCherry positive in response to LLOMe treatment, much fewer LdLPVs were CHMP2B-mCherry positive. This difference in the recruitment of these two ESCRT III components was not what we had observed in the studies of the recruitment of CHMP2B and CHMP4B to LdLPVs in infected cells (Figure 1). We interpreted these results to mean that ESCRT III recruitment to LdLPVs in infected cells is mechanistically different from the recruitment of ESCRT III molecules to damaged lysosome or lysosome-like membranes.

There is still uncertainty on the range of conditions that trigger the recruitment of ESCRT components to membranes. Membrane damage caused by rupture of endomembranes including the insertion of secretion apparatus into membranes by intracellular pathogens has been shown to be a reliable trigger for the recruitment of ESCRT components [34,38]. Our current understanding of LdLPV biology has not revealed the existence of a secretion apparatus or a process that may cause damage to the LdLPV and by so doing activate recruitment of ESCRT components. Instead, we had reported previously on the observation that phosphoinositides, specifically PI(3,4)P2 are displayed on LPVs [48]. Informed by studies that have identified PI(3,4)P2 and other phosphoinositides as membrane anchors for the recruitment of ESCRT components [42,44] we affirmed that LdLPVs do indeed display PI(3,4)P2, and then observed that treatment with LY294002 led to reduced display of this lipid and also reduced recruitment of CHMP4B to LdLPVs. The experiments with LY294002 were challenging because prolonged incubation of infected cells in this drug (greater than 8hrs) resulted in death of parasites and clearance from infected cells. We did not determine what was the major cause of cell death in the presence of the drug; however, we had reported previously that the display of phosphoinositides on LPVs was necessary for sustained activation of Akt, which rendered infected cells refractory to inducers of macrophage activation that may lead to parasite death [48]. Future studies will attempt to identify the specific ESCRT molecule(s) that are initially recruited to the LdLPV and the conditions that lead to non-canonical activation of the ESCRT machinery. It is expected that there are additional signals that are elaborated by this parasite that instruct the machinery to execute the fission and scission of LPVs, which is a phenomenon that does not occur with other pathogen containing compartments.

The membrane anchor for ESCRT components is an important issue that has been considered in several pathogen-host situations. In the budding of HIV, the Gag protein of the virus was shown to be the primary viral molecule that interacts with ESCRT [55,56]. Following from such studies small molecules that selectively block the HIV Gag – ALIX interaction have been proposed as potential viable inhibitors for the spread of HIV ([57]). In *Toxoplasma* infections, proximity labeling studies revealed interactions between ALIX (PDCD6IP) and so called GRA proteins discharged from the parasites’ Rhoptry organelle that are displayed on the parasite vacuole ([41]). Other studies had implicated the interactions of TSG101 in interactions with other GRA proteins that were necessary for nutrient acquisition into the parasitophorous vacuole ([40]). There are several interactions between ESCRT components and parasitophorous vacuoles that may satisfy different functions.

Based on our observations, we suggest that ESCRT components are recruited constitutively to LdLPVs where they are anchored to phosphoinositides that are displayed on the LdLPV membrane. The parasite likely plays some role in the synthesis of the phosphoinositide that are displayed on the LdLPV membrane. It was not determined which ESCRT molecule(s) initially anchors to the LdLPV. We do not exclude the likelihood that other ESCRT components are recruited prior to the recruitment of ALIX. However, the recruitment of ALIX is followed by the recruitment of CHMP4b and CHMP2b. Subsequent activation of the ESCRT machinery which is characterized by the aggregation and/or nucleation of ESCRT III occurs at the time of parasite division. In some images, we observed CHMP4b most prominently, at the bridge between two LdLPVs presumably at the point of scission (Figure 1 and supplemental Figure 1b). In the graphical abstract we highlight the observation that when ALIX levels are limiting, LdLPVs are large and can harbor 4 - 10 parasites (Figure 7). Evidently, there is a defect in the assembly of the ESCRT machinery and the inability to execute scission of the LdLPV. What could be the signal(s) for LdLPV scission? Could it be like signals for cytokinesis? Alternatelively, could it be like signal(s) for the initiation of viral budding where the interactions of ALIX and the Gag protein have been shown to be critical for the progress of that process? Future studies will likely address these questions.

**Figure 7.**
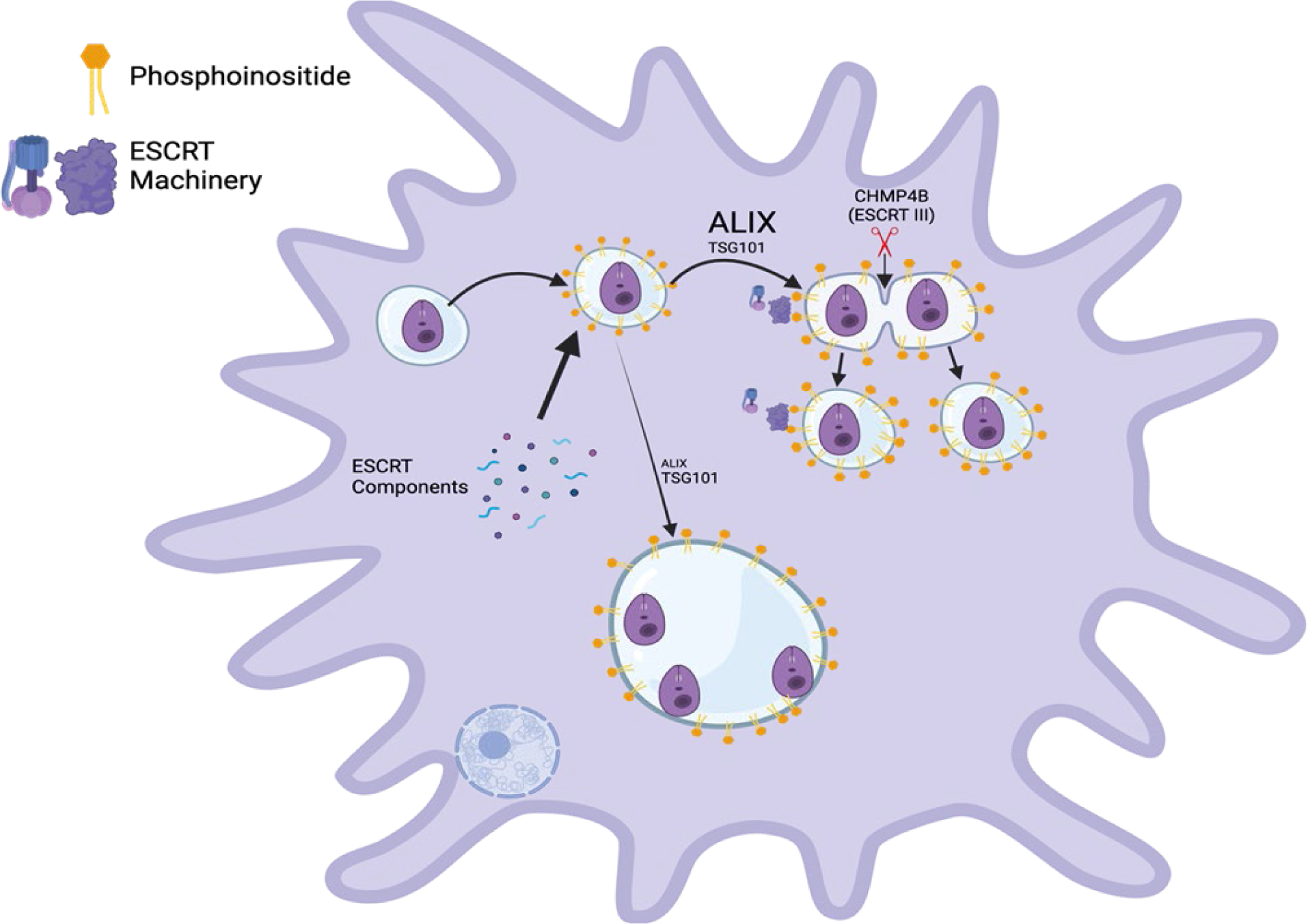
Graphical summary: LdLPVs display phosphoinositides that help anchor members of the ESCRT machinery to LdLPVs. A critical role is exercised by ALIX for activation of the ESCRT machinery to complete the scission of LdLPVs. When ALIX expression is limited LdLPV fission is defective and so too is parasite replication. This results in a reduction in the parasite burden.

## Materials and Methods

### Parasite Culture

*L. donovani* (MHOM/S.D./62/1S-CL2_D_) was obtained from Dr. Nakhasi’s lab (FDA) and cultivated in M199 media (Sigma) containing 20% FBS, 0.1 mM Adenosine, 0.1 mg/mL folic acid, 2 mM glutamine, 25 mM HEPES, 100 units/mL penicillin/-100 µg/mL streptomycin 15140122 (Gibco), 1X BME vitamins (Sigma), and 1 mg/mL sodium bicarbonate with pH 6.8 at 26°C.

### Mammalian Cell Culture

RAW264.7 macrophages were obtained from ATCC (TIB-71) and maintained in DMEM (Corning) supplemented with 10% FBS and 100 units/mL penicillin/-100 µg/mL streptomycin 15140122 (Gibco). Cells were kept in complete DMEM at 37°C in a humidified atmosphere incubator containing 5% CO_2_.

### Macrophage Infections

RAW264.7 macrophages were plated in culture dish containing sterile glass coverslips and incubated overnight at 37°C with 5% CO_2._ For infection, metacyclic promastigotes were selected from late stationary stage parasites using peanut agglutination (PNA) according to an established protocol described previously ([58]). Briefly, 4-day old cultures of *L. donovani* parasites were washed twice and resuspended in incomplete DMEM at a concentration of 2×10^8^ parasites/mL. PNA was then added to the parasites at a final concentration of 50 µg/mL and incubated at room temperature for 15 mins. The parasites were then centrifuged at 200x*g* for 5 mins to pellet agglutinated parasites. The supernatant was then collected and the PNA-metacyclic parasites were washed twice, resuspended in complete DMEM, and counted for infection. Parasites were then added to macrophage dishes at a ratio of 5:1 or 10:1 MOI (parasites:macrophages) then incubated 37°C with 5% CO_2._ Infections were stopped at 24, 48 or 72 hours post infection by placing coverslips in 4% paraformaldehyde for 20 minutes for fixation at room temperature. After fixation, cells were ready for the immunofluorescence assay. For experiments to evaluate recruitment of proteins of interest to generic phagosomes, latex beads L0280 (SIGMA-ALDRICH) were incubated with macrophages at an MOI equal to parasite infections

For PI3K inhibition, 50μM LY294002 (Cell Signaling) was added to cultures after 24 hrs infection. Coverslips from the infection were fixed after 8 and 24 hrs of incubation with the drug. Some infected cultures received DMSO (vehicle) only.

### Generation of Transfectants and knock downs in RAW264.7 cells

To develop cells that express fluorophore-tagged variants of ESCRT molecules, we acquired the following plasmids. pLNCX2-mCherry-CHMP2B was a gift from Sanford Simon (Addgene plasmid # 115331), pLNCX2-mCherry-CHMP4B was a gift from Sanford Simon (Addgene plasmid # 116923), pLNCX2-mCherry-VPS4A was a gift from Sanford Simon (Addgene plasmid # 115334), pEGFP-VSP4-E228Q was a gift from Wesley Sundquist (Addgene plasmid # 80351), pLNCX2-mEGFP-TSG10 was a gift from Sanford Simon (Addgene plasmid # 116925). To approximately 1X10^7^ RAW264.7 cells, 10ug plasmids were introduced by nucleofection using reagents from Mirus Bio (AMAXA Nucleofector II). For the most part, cells were used in experiments after transient transfections. 24 hours after nucleofection, cells were infected with *L. donovani* parasites or incubated with latex beads as described above. To develop knock downs cell lines, the following plasmids were obtained: TSG101 shRNA Plasmid sc-36753-SH (Santa Cruz), Alix shRNA Plasmid sc-60150-SH (Santa Cruz) and Control shRNA Plasmid sc-108060-SH (Santa Cruz). After nucleofection (AMAXA Nucleofector II), cells were seeded in 24 well plates for selection of oligoclonal lines. After 24 hours, the culture medium was replaced with medium containing 6ug/ml of puromycin (it was predetermined that the RAW264.7 wild type cells were killed by 4ug/ml puromycin). Colonies were selected from the 24 well plates and expanded into 6 well plates. Oligoclonal lines in 6 well plates were expanded for downstream application and experimentation. Knock down of specific proteins was validated by Western Blotting as described below.

### LDH Cytotoxicity and MTT Proliferation Assays on Protein Knocked Down Cells

To test if cells that were ALIX or TSG101 knocked down were viable, we performed LDH (Lactate dehydrogenase) cytotoxicity assay with the CK12 (DOJINDO) cytotoxicity kit. 5000 cells were seeded in 96 well plates in 4 wells per cell line and separate 96 well plates for each time point. Then, assessment of LDH release was followed per kit instructions, and cytotoxicity was determined using LDH release from cells that were transfected with the control plasmid as reference. For MTT proliferation assay, 2000 cells were seeded in 96 well plates in 4 wells per cell line and separate 96 well plates for each time point. Proliferation was assessed using the MTT cell proliferation kit AR1156 (BOSTER), and kit protocol was followed for each collected timepoint.

### Immunofluorescence Assay (IFA) and Fluorescence Microscopy

Glass coverslips with infected macrophages were fixed in 4% paraformaldehyde (PFA) at the indicated times post infection. PFA was washed out after 20 minutes of incubation, and then washed with PBS twice. Coverslips were left in PBS until all timepoints of infection were collected for downstream application. Coverslips were then processed in immunofluorescence assays (IFAs) to visualize the distribution of LAMP1 to delineate parasite vacuoles, PI(3,4)P2 for phosphoinositides. Anti-LAMP1 1D4B (DSBH), and/or anti PI(3,4)P2 Z-P034 (Echelon Biosciences) primary antibodies were diluted at a 1:50 and 1:100 concentrations, respectively. Chicken anti-rat AF594 conjugated A21471 (Invitrogen), goat anti-rat AF488 conjugated A11006 (Invitrogen) or chicken anti-mouse FITC conjugated NB120-6810 (NOVUS) secondary antibodies were diluted in PBS at a 1:200 concentration. Briefly, coverslips were washed in 1XPBS, then incubated for 10 minutes in 50mM ammonium chloride to quench remaining PFA. Cells were permeabilized in 0.5% Triton-X-100 in 1XPBS for 15 minutes and blocked in 5% bovine serum albumin (BSA). Incubation in primary antibodies diluted in 1XPBS with 1% BSA was performed for 1 hour at room temperature. After washing in 1XPBS, coverslips were incubated in AlexaFluor/FITC conjugated secondary antibodies for 30 minutes at room temperature. After washing, coverslips were mounted on glass slides using the ProQ diamond mounting agent, which is supplemented with 4’,6-diamidino-2-phenylindole (DAPI) stain. IFAs were visualized and captured using BZ-X810 (Keyence, Osaka, Japan) All-in-One Fluorescence Microscope at 100x oil immersion objective magnification or Zeiss Axio Observer Z1/7; 63× water immersion objective magnification. Images were captured with an Axiocam 503 controlled by Zen Acquisition software. Scored LdLPVs were delimited by Lamp1 reactivity that contained at least one parasite nucleus. The percentage of infected macrophages and the average number of parasites was determined by counting at least 100 macrophages per coverslip for LdLPVs or latex bead phagosomes. Counts were done in duplicate coverslips for at least 3 independent experiments.

### Image Analysis

Images were analyzed using the BZ-X800 analyzer for pictures taken in the BZ-X810 fluorescent microscope, or Zen Blue 2.6 for pictures taken in the Zeiss Axio observer Z1/7. Z stack projections were used for specific image analysis of colocalizing proteins. Using Fiji/ImageJ, a mask was created around parasite vacuoles or latex bead phagosomes determined to be regions of interest (ROI). Then, the BIOP JAcOP colocalization of signals plug-in was used to determine if signals of proteins of interest colocalized in the ROI previously selected using the masking option. Lamp1 reactivity was used to delineate the vacuole of the parasite or latex bead phagosome; colocalization of Lamp1 with ESCRT proteins of interest was performed. If the Pearson’s correlation coefficient yielded by JACoP met a specific threshold for the masked region, then it was counted as positive. At least 100 parasite vacuoles or latex bead phagosomes were counted per coverslip, per treatment and timepoint. This analysis was applied for each experimental scheme.

### LLOMe Pulse/Chase Assay

To determine L-Leucyl-L-Leucine methyl ester (LLOMe) hydrobromide HY-129905A (MedChemExpress) impact on parasite vacuoles, we prepared a stock solution of 100mM using dimethyl sulfoxide (DMSO) as a solvent. LLOMe was diluted with complete DMEM at 5mM and 1mM concentrations and added to infected macrophages 48 hours post infection. Infected cells were stimulated for 30 minutes with each LLOMe concentration and an equal amount of DMSO was added to complete DMEM for vehicle control and added simultaneously on separate vessels. After 30-minute exposure, LLOMe was washed out twice with complete DMEM, and coverslips were collected post LLOMe stimuli at 0 minutes, 10 minutes, 30 minutes and 60 minutes.

### Parasite Vacuole Measurement, Parasites per Vacuole and Parasites per Macrophage Counts

Z-Stack images of infected cells on coverslips were acquired as described above. Z-Stack heights were selected to cover the entire host cell nucleus and any PVs being measured to ensure that the widest point of the host cell nucleus and PV were captured. Using the “Insert” scale tool of the BZ-X800 analyzer, diameter of parasite vacuoles was measured. At least 100 parasite vacuole sizes were measured per coverslip, per treatment and timepoint. For parasite counts per macrophage, at least 100 macrophages were counted, per coverslip, per treatment and timepoint. Parasites per vacuole were counted using DAPI parasite nuclei, and Lamp1 reactivity for parasite vacuole reference.

### Western Blotting

Cells were lysed in RIPA buffer #89900 (Thermo Scientific) and protein concentration was measured by BCA assay #23225 (Thermo Scientific). Aliquots with equal protein (ug) amounts were suspended in SDS PAGE loading buffer and run at 140V for 1hr. Protein was transferred onto a nitrocellulose membrane 10600006 (GE Healthcare) or polyvinylidene difluoride membrane IPVH15150 (Immobilon) with 20V for 10hr in 4°C. Primary antibodies included anti-Alix sc-53540 (Santa Cruz), anti-TSG101 PA5-81094 (Invitrogen) and anti-Beta Actin sc-47778 (Santa Cruz). Secondary HRP conjugated antibodies included Goat anti-mouse IgG 31430 (Invitrogen) and Chicken anti-Rabbit IgG A15993 (Invitrogen). Antibody dilutions were prepared in 1X TBST (Tris buffered saline Tween 20) with 5% BSA for primary antibodies, and 5% non-fat milk for secondary antibodies. Primary antibodies were diluted at a 1:1000 concentration, and secondary antibodies at a 1:2000 concentration. Membranes were blocked with 5% non-fat milk for 1hr then incubated with primary antibody overnight at 4°C in constant rocking. After washing with TBST 3 times for 10 minutes each, secondary antibody incubation was performed for 1 hour. Proteins of interest were then visualized using chemiluminescence on the Invitrogen iBright imaging system (Thermo Fisher Scientific). Protein quantification was performed using FIJI/imageJ’s densitometric analysis tool.

### Statistical Analysis of Data

After data acquisition and data collected by colocalization of proteins of interest, parasite vacuole sizes, parasite infectivity, parasites per vacuole and parasites per macrophage were graphed using GraphPad Prism 8. Data was first imported, then a one-way ANOVA or unpaired Student T Test was performed, and statistical significance was determined by *post hoc* Tukey’s honest significant difference test in the case of ANOVA. Similarly, densitometry data for western blot quantification of protein levels was acquired from FIJI/ImageJ as described above, then GraphPad Prism 8 was used to generate bar graphs and perform unpaired student T Test. Significance was determined by a p-values of *p<0.05, **p<0.01, ***p<0.001, ****p<0.0001

## Supplemental Figures

**Figure S1.**
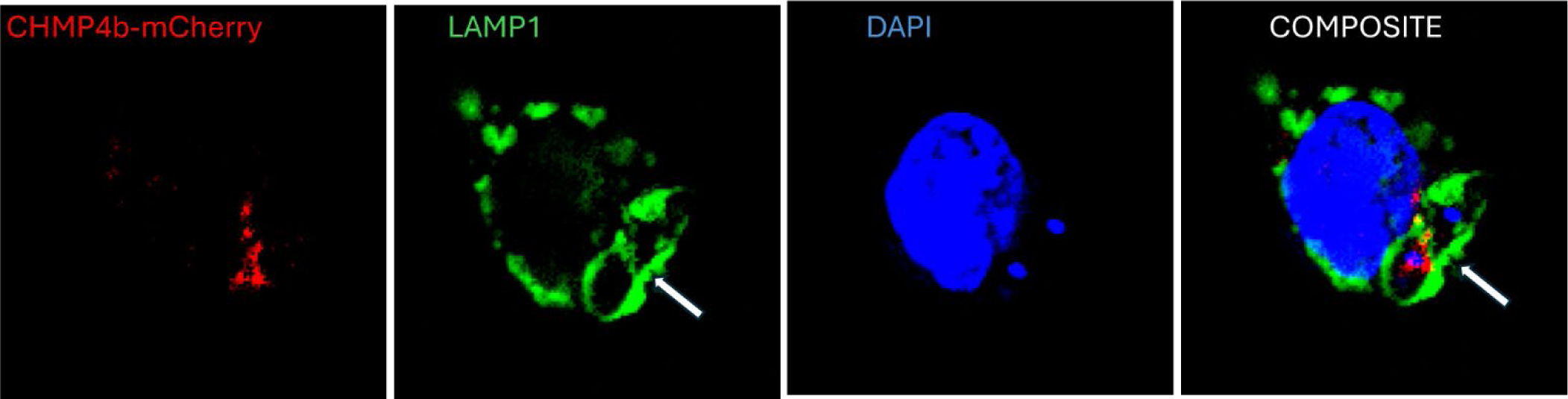
CHMP4B is recruited to LdLPVs. Representative image that shows CHMP4B in the middle of two LdLPVs.

**Figure S2.**
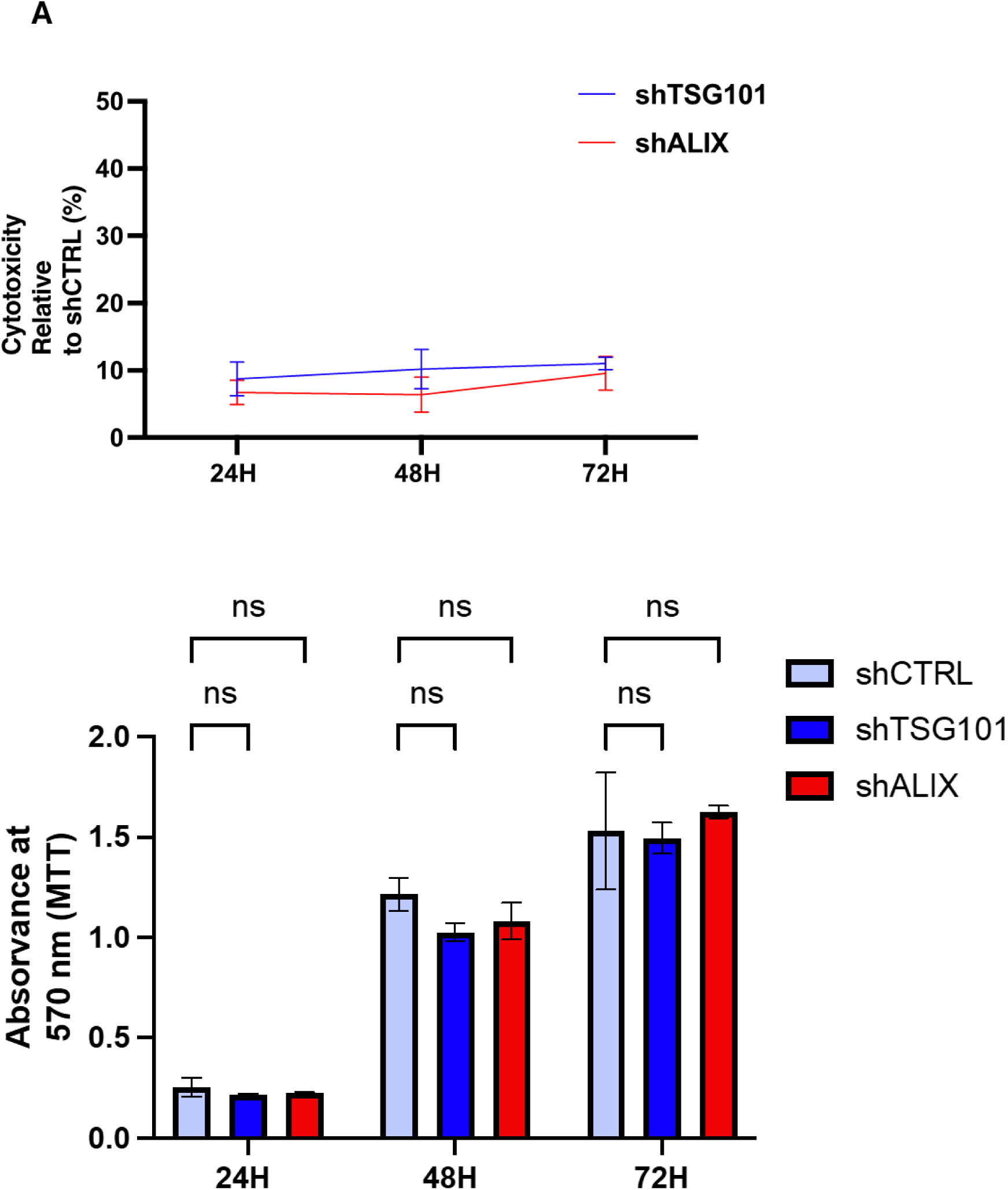
Neither TSG101 knock down nor ALIX knock down is cytotoxic to cells Cytotoxicity was determined by monitoring LDH release from cells knock down oligoclonal lines and from sCTRL (A). For MTT proliferation assay, cells were seeded in 96 well plates in 4 wells per cell line and separate 96 well plates for each time point. Proliferation was assessed using the MTT cell proliferation kit AR1156 (BOSTER) (B). Results are from 2 experiments.

